# Development and validation of an open-source, disposable, 3D-printed *in vitro* environmental exposure system for Transwell^®^ culture inserts

**DOI:** 10.1101/2020.10.27.358168

**Authors:** Abiram Chandiramohan, Mohammedhossein Dabaghi, Jennifer A. Aguiar, Nicholas Tiessen, Mary Stewart, Quynh T. Cao, Jenny P. Nguyen, Nima Makhdami, Gerard Cox, Andrew C. Doxey, Jeremy A. Hirota

**Affiliations:** Firestone Institute for Respiratory Health – Division of Respirology, Department of Medicine, McMaster University, Hamilton, ON, L8N 4A6; Department of Biology, University of Waterloo, Waterloo, ON, Canada N2L 3G1; McMaster Immunology Research Centre, McMaster University, Hamilton, ON, Canada, L8S 4K1; Division of Respiratory Medicine, Department of Medicine, University of British Columbia, Vancouver, BC, Canada, V6H 3Z6

## Abstract

Accessible *in vitro* models recapitulating the human airway that are amenable to study whole cannabis smoke exposure are needed for immunological and toxicological studies that inform public health policy and recreational cannabis use. In the present study, we developed and validated a novel 3D printed *In Vitro* Exposure System (IVES) that can be directly applied to study the effect of cannabis smoke exposure on primary human bronchial epithelial cells.

Using commercially available design software and a 3D printer, we designed a four-chamber Transwell^®^ insert holder for exposures to whole smoke. Software was used to model gas distribution, concentration gradients, velocity profile and shear stress within IVES. Following simulations, primary human bronchial epithelial cells cultured at air-liquid interface on Transwell^®^ inserts were exposed to whole cannabis smoke. Following 24 hours, outcome measurements included cell morphology, epithelial barrier function, lactate dehydrogenase (LDH) levels, cytokine and gene expression.

Whole smoke delivered through IVES possesses velocity profiles consistent with uniform gas distribution across the four chambers and complete mixing. Airflow velocity ranged between 1.0-1.5 μm s^−1^ and generated low shear stresses (<< 1 Pa). Human airway epithelial cells exposed to cannabis smoke using IVES showed changes in cell morphology and disruption of barrier function without significant cytotoxicity. Cannabis smoke elevated IL-1 family cytokines and elevated CYP1A1 and CYP1B1 expression relative to control.

IVES represents an accessible, open-source, exposure system that can be used to model varying types of cannabis smoke exposures with human airway epithelial cells grown under air-liquid interface culture conditions.

## Introduction

The lung is responsible for gas exchange between the outside world and the human body. During ventilation, the lungs are in contact with the external environment and exposed to air that may contain a variety of pathogens including allergens, viruses, bacteria, and air pollutants, including organic combustion byproducts (Huff, Carlsten & Hirota, 2019). The pseudostratified airway epithelial cells that line the lungs have evolved to mitigate the risks from external insults, providing a tight physical and immunological barrier (Hirota & Knight, 2012; Iwasaki et al., 2017; Whitsett & Alenghat, 2015). It contributes to coordinated immune responses including mucus production and airway surface lining fluid secretion, cytokines and chemokines secretion for local and systemic immune cell recruitment, and antimicrobial protein secretion in response to environmental assaults (Ganesan et al., 2013; Bakshani et al., 2018). Airway epithelium dysfunction has been strongly implicated in the pathogenesis of many airway diseases including asthma, Chronic Obstructive Pulmonary Disease (COPD), and pulmonary fibrosis (Kicic et al., 2006; Steiling et al., 2013; Xu et al., 2016). Importantly, environmental insults capable of damaging the airway epithelium are also risk factors for these same respiratory diseases (Broekema et al., 2009; Tamashiro et al., 2009; Thorley & Tetley, 2007; Walters et al., 2014).

Tobacco smoking, a direct environmental insult to the airway epithelium, remains common on a global scale despite well described effects on respiratory health (Amatngalim et al., 2016; Cantin, 2010; Mathis et al., 2013; Schamberger et al., 2015; Thun et al., 2002). Similar to tobacco smoke, cannabis smoking is also a direct environmental attack to the lungs. As a result of progressive legalization on a global scale, cannabis consumption has been on the rise, where 90% of consumers prefer smoking as a route of delivery (World Drug Report, 2019; Canadian Cannabis Survey, 2018). Tobacco and cannabis smoke exposure induce similar clinical features such that both environmental exposures are associated with increased prevalence of coughing, wheezing, chest tightness, and risk for developing COPD (Moir et al., 2008; Tashkin et al., 1987; Wu et al., 1988). Tobacco and cannabis smoke exposure also present divergent clinical features, with tobacco smokers more likely to develop lung cancer relative to cannabis smokers (Melamede, 2005). This observed divergence may be due to the presence of phytocannabinoids with anti-inflammatory properties unique to cannabis (Kaplan et al., 2008; Nagarkatti et al., 2009; Tashkin et al., 1975; Vachon et al., 1973). The uncertainty surrounding the health effects of cannabis smoke and discrepancies with the known negative impacts of tobacco smoke warrants further investigation to inform government policies, recreational practices and cultivation strategies in an era of broadening acceptance for legal and open markets.

*In vitro* smoke exposure models using human airway epithelial cells have been crucial for tobacco smoke research and will likely be important for cannabis smoke research (St-Laurent et al., 2009; Wan et al., 2009). A proportion of available smoke exposure research, primarily with tobacco, has been established through submerged monolayer culture designs in which smoke extract has been used to expose cells (Ji et al., 2017; Jamieson et al., 2019; Gellner et al., 2016). Our group has recently applied these methods to Calu-3 cells with cannabis smoke extract exposure and observed an induction of a pro-inflammatory cytokine response and suppression of antiviral cytokines (Aguiar et al., 2019; Huff et al., 2020). Smoke extract exposures do not entirely reflect whole smoke exposure as the latter generates heat and water insoluble hydrocarbon combustion products (Lange, 2007; Moir et al., 2008; Schwartz, 2017; Tashkin et al., 1991). Additionally, submerged monolayer cell culture designs do not entirely reflect the *in situ* airway environment where a pseudostratified airway epithelium is exposed to inhaled air on an apical side while attached to a basement membrane on the basal side. To more accurately model the *in situ* environment, airway epithelial cells can be cultured on Transwell^®^ inserts under air-liquid interface (ALI) culture conditions to create a pseudostratified tissue structure (Azzopardi et al., 2015; Jiang et al., 2018). Transwell inserts of airway epithelial cells under ALI can be used for whole smoke exposures using advanced systems that model inhalation and exhalation patterns of humans (Azzopardi et al., 2015; Jiang et al., 2018). The strengths and limitations of popular smoke generating machines have been characterized extensively (Aufderheide & Mohr, 1999; Deschl et al., 2011; Ritter et al., 2001; Thorne & Adamson, 2013). Commercially available smoking machines offer high throughput designs and are able to accommodate multiple cigarettes or cell culture plates that can be exposed with independent syringes, allowing for control over the parameters of each exposure under automated conditions (Thorne & Adamson, 2013). Some devices are directly amenable to multiple exposures beyond smoke, including environmental toxins, gases, therapeutic aerosols, aerosolized pathogens, and other volatile compounds (Aufderheide & Mohr, 1999; Deschl et al., 2011; Ritter et al., 2001; Thorne & Adamson, 2013). The throughput and automation benefits of these environmental exposure systems are offset by some important limitations (Adamson et al., 2011; Aufderheide et al., 2003; D. W. Bom bick, 1997; Scian et al., 2009; Thorne & Adamson, 2013). From an operations perspective, these are capital intensive closed systems that cannot be customized, limiting accessibility to specialized research facilities and field participants (Thorne & Adamson, 2013). From a technical perspective, limitations may include potential air and smoke mixing inefficiencies, aging of smoke in mixing vessels, and turbulence that is inconsistent with lung physiology (Thorne & Adamson, 2013). Collectively, the development and validation of a low-cost *in vitro* environmental exposure system for Transwell^®^ culture inserts of ALI cultured airway epithelial cells will broaden the ability of researchers to perform essential research related to cannabis smoke exposure and lung health.

To address existing constraints with environmental exposure systems and related research, we propose that additive manufacturing, such as 3D printing, can function as a disruptive solution to create an open-source, disposable, *in vitro* exposure system that is widely accessible without requiring specialized environmental exposure infrastructure. Additive manufacturing has been utilized in various fields to realize complex 3D constructs with a resolution at the micron level. In contrast to conventional manufacturing technologies, additive manufacturing is accessible and independent of specialized technologies or personnel, with widely available commercial 3D printers able to rapidly generate functional designs based on computer aided design (CAD) files. Additive manufacturing technologies have accelerated prototyping steps while reducing costs, effectively enabling researchers with limited design training to optimize novel designs independent of historical manufacturing constraints. Importantly, commercially available and capital accessible additive manufacturing technologies are sufficient for the medium throughput production that is required at a standard research lab scale.

In the present study, we develop and validate an open-source, disposable, 3D-printed *in vitro* environmental exposure system for Transwell^®^ culture inserts and apply to study the effect of whole cannabis smoke on primary human bronchial epithelial cells (HBECs). Our *In Vitro* Exposure System (IVES) is widely accessible due to the open-source CAD file, amenability to commercially available 3D printers, and design for widely used Transwell^®^ culture inserts. We validate our model with cannabis smoke laying a foundation for additional modalities including tobacco smoke and vaping products. IVES also gives the capacity for four transwell inserts to be exposed to the same environmental exposure, providing medium throughput while reducing variability that may occur with independent exposures for separate transwells. Finally, IVES is amenable to dilution of smoke with room air for adaptation to concentration-response studies. Using cannabis smoke as an exposure of current public health importance, we validate IVES with primary HBECs and demonstrate an impact of the exposure on epithelial barrier function, IL-1 family cytokine expression and expression of genes involved in cellular detoxification.

## Materials and Methods System

### System Design

Autodesk Inventor software (2018, Autodesk, CA, USA) was used for all designs. The IVES units were 3D-printed with a FormLabs Form 2 printer using clear resin (RS-F2-GPCL-04, MA, USA). IVES contains four Transwell exposure chambers, two inlets, four outlets, and four chamber caps (**Figure 1**). Each inlet is distributed to each of the four exposure chambers. Inlet diameter was optimized at 3 mm to minimize obstruction with organic combustion byproducts. Exposure chamber size was designed based on the dimension of a Transwell^®^ insert for 24 well-plate. In addition, the location of the inlet and outlet of each chamber was designed such that Transwell^®^ inserts would experience indirect exposures of turbulent air. Each chamber had an outlet for exhaust from the chamber. Threaded caps for the exposure chambers created seals that eliminated exhaust through this path.

**Figure 1:**
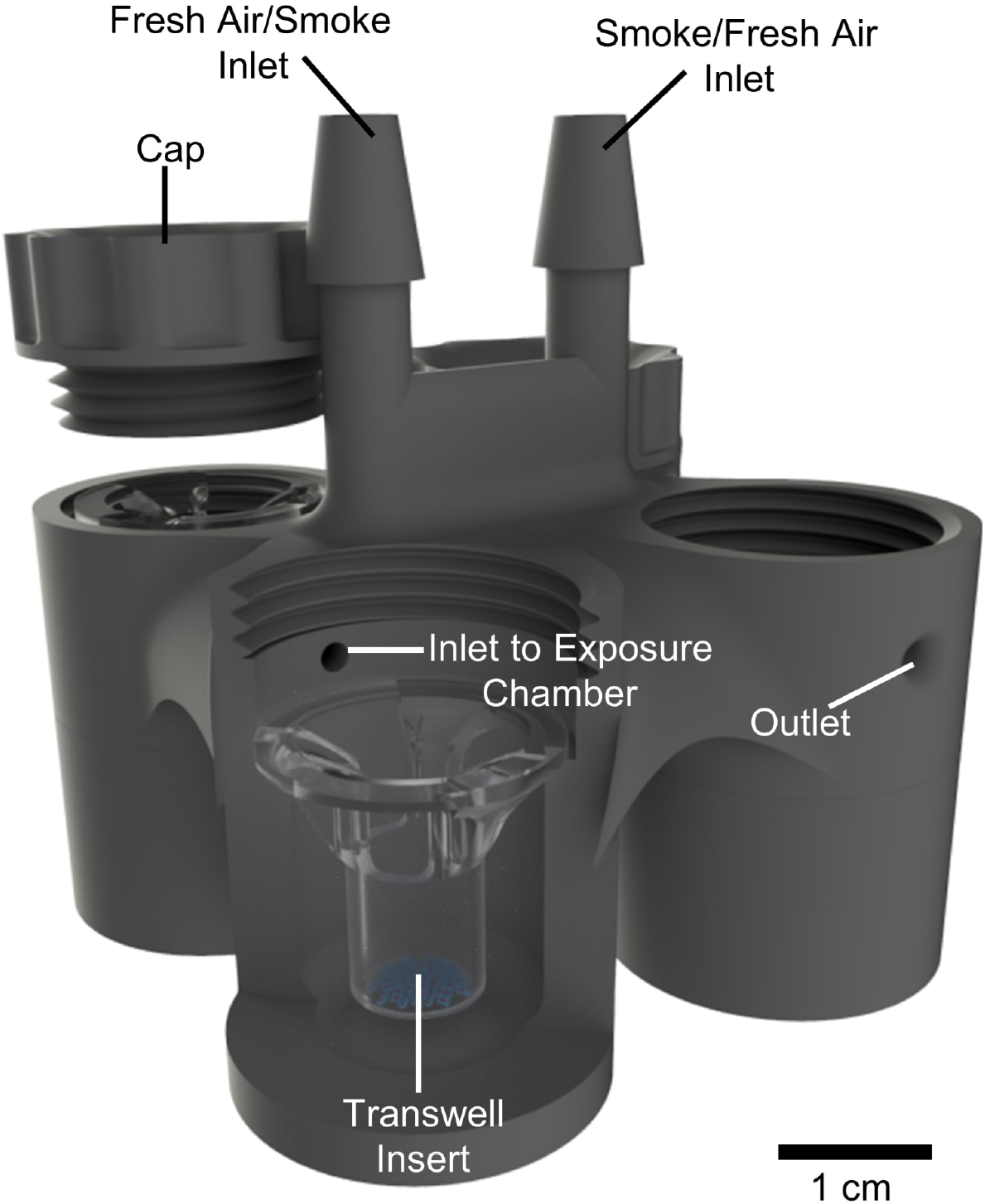
3D schematic view of the *in vitro* exposure system (IVES). As shown, air or exposure (e.g. smoke) delivered to the inlet is equally distributed across the four exposure chambers which house the Transwell^®^ inserts. Circulated air or exposure (e.g. smoke) exits passively through the outlet in each exposure chamber.

### Fluid Dynamic Modeling

During design phases, a quantitative simulation of fluid dynamics was performed using COMSOL Multiphysics software to model the gas distribution and concentration gradients in the IVES unit. To model the velocity profile and shear stress, the entire fluidic path from the merged inlet to the chambers and the outlets were included. The main purpose of this simulation was to ensure that the gas was equally distributed among all chambers with minimal shear stress experienced at the surface of Transwell^®^ inserts. Air was used as the gas of interest in this part of the simulation, under the assumption that air was behaving as a Newtonian fluid. The governing equations used in the simulation as follows:

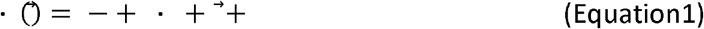

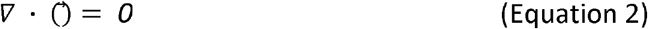

Where ρ is the air viscosity, 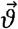 represent the velocity vector field, p shows the pressure, 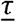 express the stress tensor field, ^→^ is the gravitational force, and shows for any possible external source term. The acceleration forces, including both the local and the conventional forces, are expressed on the left side of Equation 1, and the forces created by pressure gradient and viscosity are described on the right side of the equation. A no-slip condition on the walls was assumed for the entire IVES unit, with the model assuming no back-flow. A gas flow rate of 7 mL s^−1^ was used as the boundary condition and the gas flow direction was set to be normal to the inlet and outlets. Equation 2 expresses the mass conservation for the exposure unit. In the simulation, initial values were set to zero. Then, the simulation was studied under a steady-state condition.

### Cell Culture

All studies using human cells were approved by the Hamilton Integrated Research Ethics Board (HiREB Approval: 5305T). Primary HBECs were isolated from a bronchial brushing from consented subjects undergoing routine clinical procedures, and plated into T25 flasks containing Pneumacult™ Ex-Plus Basal Media (StemCell Technologies; Catalog Number 05040) with PneumaCult™ Ex-Plus 50x Supplement, 0.01% hydrocortisone stock solution (StemCell Technologies; Catalog Number 7925) and 1% antibiotic-antimycotic (Thermofisher; Catalog Number 15240062). Once cultures achieved ~80% confluence, cells were passaged at a density of 50,000 - 100,000 cells/insert into polyester Transwell^®^ inserts for exposure experiments. Cells were fed with 200μl and 750μl of Pneumacult™ Ex-Plus Basal Media in the apical and basal compartments, respectively. Once cultures reached 100% confluency, apical media was removed and culture inserts were fed from the basal side to bring cultures to ALI (Day 0 of ALI). Following 24h, culture inserts were fed from the basal compartment with 750μl of PneumaCult™ ALI Basal Medium (StemCell Technologies; Catalog Number 05001) with PneumaCult™-ALI 10x Supplement, PneumaCult™-ALI Maintenance 100x Supplement, 1% antibiotic-antimycotic, 0.5% hydrocortisone stock solution and 0.2% heparin solution (StemCell Technologies; Catalog Number 7980) to support development and differentiation of a pseudostratified epithelial culture (Day 1 of ALI). Transwell^®^ cultures were fed from the basal compartment and a phosphate buffered saline (PBS) wash was performed on the apical compartment every other day. Exposure experiments were performed between Day 14 - Day 15 of ALI culture.

### Cannabis Material and Cigarette Manufacturing

A Kentucky Research Grade Cigarette (Lot:3R4f) contains ~0.7g of dried tobacco leaves and has been used extensively in *in vitro* studies. Using research grade tobacco cigarettes as a reference, cannabis cigarettes with ~0.7g of dried cannabis flower (~10% THC, 0% CBD; *Purple Sun God*, Lot: 00117 (b161)) were manufactured with RAW rolling papers and cardboard filters. Cannabis was purchased from the Ontario Cannabis Store with a Health Canada approved research license.

### Epithelial Barrier Function Assessment

Transepithelial Electrical Resistance (TEER) was measured using a Millicell ERS-2 Voltohmmeter (EMD Millipore, Etobicoke, Ontario, Canada) to quantify epithelial barrier function according to manufacturer’s directions. TEER was measured prior to and 24h post exposures (air or smoke).

### Exposure Protocol

In a biosafety cabinet, 800ul of PneumaCult™ ALI media was added into each individual chamber of the IVES unit. Transwell^®^ inserts were transferred into the IVES and left in the a 37°C, 5% CO_2_ humidified incubator to equilibrate for 10 minutes prior to exposures. In a fume-hood, exposures were performed with a 50ml syringe connected to the fresh air inlet of the IVES through a PVC Tygon tube (length: 32 cm, lumen diameter: 2 mm) and a 3-way valve (**Figure 2**). Another 50ml syringe was connected to the smoke exposure inlet on the IVES through PVC Tygon tube (length: 32 cm, lumen diameter: 2 mm) and a 3-way valve. For the experiments with cannabis, a cigarette was placed into another PVC Tygon tube (length: 32 cm, lumen diameter: 8 mm), with the opening sealed with parafilm and connected to the 3-way valve. The dose was administered according to a modified version of the Foltin Puff Procedure (Abdallah et al., 2018; Chait et al., 1988; Foltin et al., 1987; Ramesh et al., 2013; Wilsey et al., 2013, 2016). This procedure, optimized for our model, consisted of 1 puff of smoke separated by 3 puffs of fresh room air to mimic the behavior of human smoking patterns. Each puff of smoke or room air was 35ml in volume and was perfused over the Transwell^®^ insert at a rate of 7 ml/second. Between each puff, the cells were undisturbed for 10 seconds. Each experimental group received one cigarette smoked to completion using this regimen. Control conditions received as many puffs of room air as the corresponding experimental condition received smoke under the same regimen. Following exposure, inserts were immediately transferred to a new plate with 600ul of PneumaCult™-ALI media in the basal compartment. Plates were then transferred to a 37°C, 5% CO_2_ humidified incubator. Twenty-four hours later, basal and apical media was collected, spun down at 7500 x g/4°C for 15 minutes and supernatants were stored at −80°C for subsequent quantification of cytokines and assessment of cytotoxicity.

**Figure 2:**
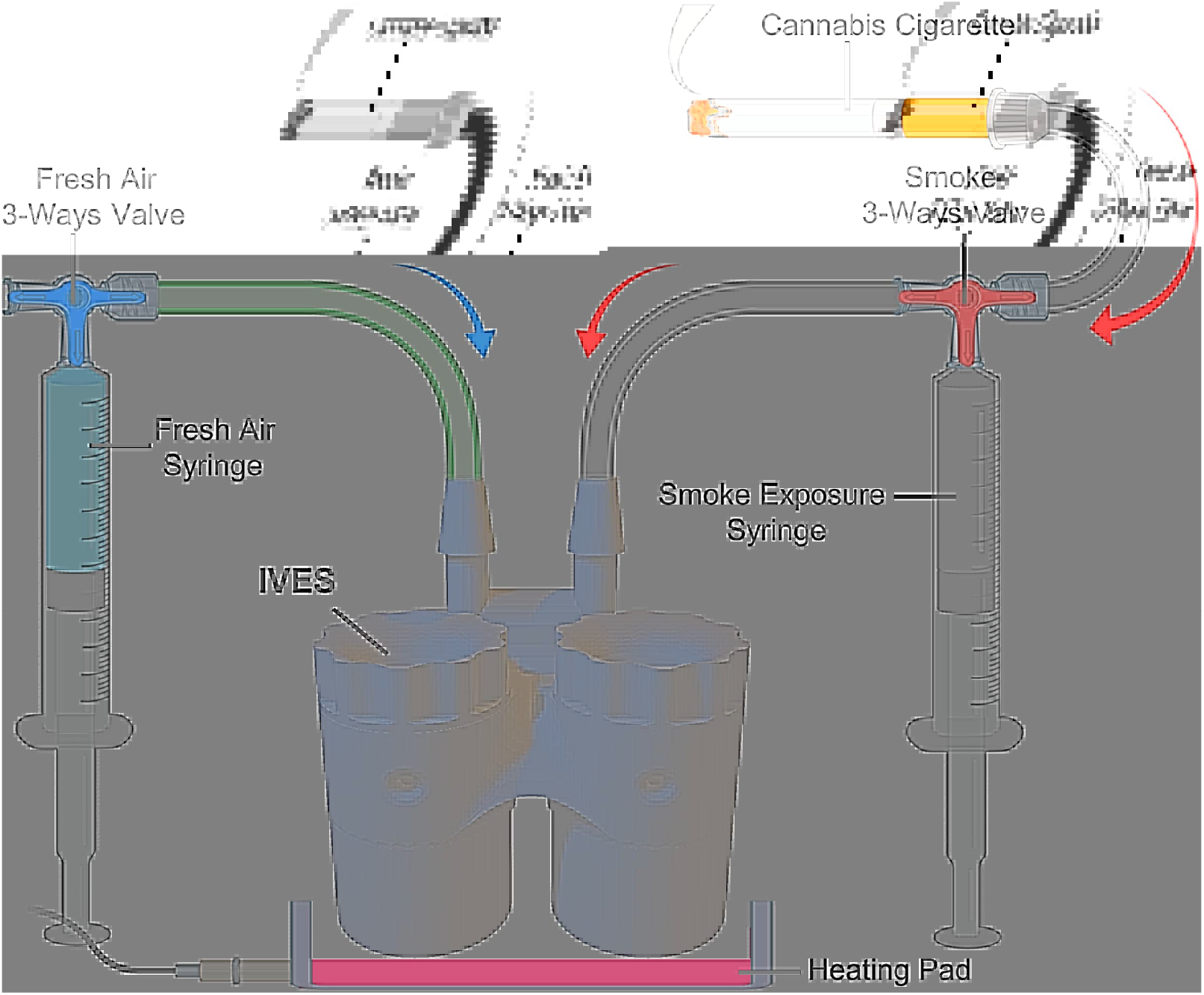
A schematic depicting the IVES connected to air and smoke sources. A 3-way valve connects the cannabis cigarette to IVES through a 50ml syringe. Another 3-way valve connects room air to IVES. Smoke is drawn through the smoke exposure syringe and expelled with predetermined rate and volume into IVES. Room air is introduced with the fresh air syringe in a similar fashion. A heating pad positioned below IVES maintains the experimental system at 37°C (Figure generated with BioRender).

### Quantification of Cytokines

Cell culture media was collected and spun down at 7500 x g for 15 minutes at 4°C. Apical supernatants were subsequently analyzed a via 42-plex multiple cytokine array (Eve Technologies, Calgary, Alberta, Canada).

### Cell Membrane Integrity and Cytotoxicity

A CYTOQUANT™ lactate dehydrogenase (LDH) Cytotoxicity Assay kit (ThermoFisher; Catalog Number C20301) was used according to the manufacturer’s guidelines with positive controls used to indicate maximal LDH release.

### RNA Isolation, Purification and Transcriptomic Analysis

Following collection of apical and basal media 24h post-exposure, total RNA was extracted and isolated using an RNeasy Kit (Qiagen, Toronto, Ontario, Canada). Cells in each Transwell^®^ were lysed in 100ul of RLT Isolation Buffer with 1% beta-mercaptoethanol (2Me) (v/v) and stored at −80°C. Subsequently, RNA was purified using the manufacturer’s protocol and quantified via Nanodrop200. cDNA was prepared and underwent transcriptomic analysis using Clariom S Microarray chips (ThermoFisher, Mississauga, Ontario, Canada).

### Processing of raw microarray data

Raw intensity values from the Clarion S microarray experiment were imported into the R statistical language environment (version 3.6.1; R Core Team, 2019). Probe definition files were obtained from the Brainarray database (version 24) (Dai et al., 2005). The Single Channel Array Normalization (SCAN) method was used to obtain log_2_- transformed normalized expression values with the SCAN.UPC R package (version 2.26.0) (Piccolo et al., 2012), with annotation data from the Bioconductor project (version 3.9) (Huber et al., 2015).

### Statistical Analysis

All statistical tests were performed on Graphpad^®^ Prism 8 version 8.3.0 (Graphpad Headquarters, San Diego, CA). A paired t-test was performed to assess significance between the differences in epithelial barrier function between the control group and experimental group. An ANOVA followed by a Tukey’s post-hoc test was performed to assess differences in cell cytotoxicity following smoke exposure. Differences in cytokine expression were assessed using paired t-tests. P values less than 0.05 were considered statistically significant.

## Results

### IVES Fluid Dynamic Modeling

A mesh analysis was conducted to determine the finest mesh resolution. The mesh size was varied from coarse to extra fine, and the mesh size was chosen to be finer since no significant change was recognized by changing the mesh size from finer to extra fine. A defined mesh resolution was chosen to run all simulations for fluid dynamic modeling (**Figure 3**) with a 3D view of velocity streamlines presented (**Figure 3a** to **Figure 3d**). Quantitative values for the velocity profile were consistent with a uniform gas distribution with complete mixing (streamlines) (**Figure 3e**). In addition, the velocity profile (**Figure 3f**) and the shear stress profile (**Figure 3g**) at the Transwell^®^ insert growth area location were simulated, investigating the impact of the exposure to the cells growing on this surface. Airflow velocity at an approximation of cell location was 1.0-1.5 μm s^−1^ and generated shear stresses << 1 Pa.

**Figure 3:**
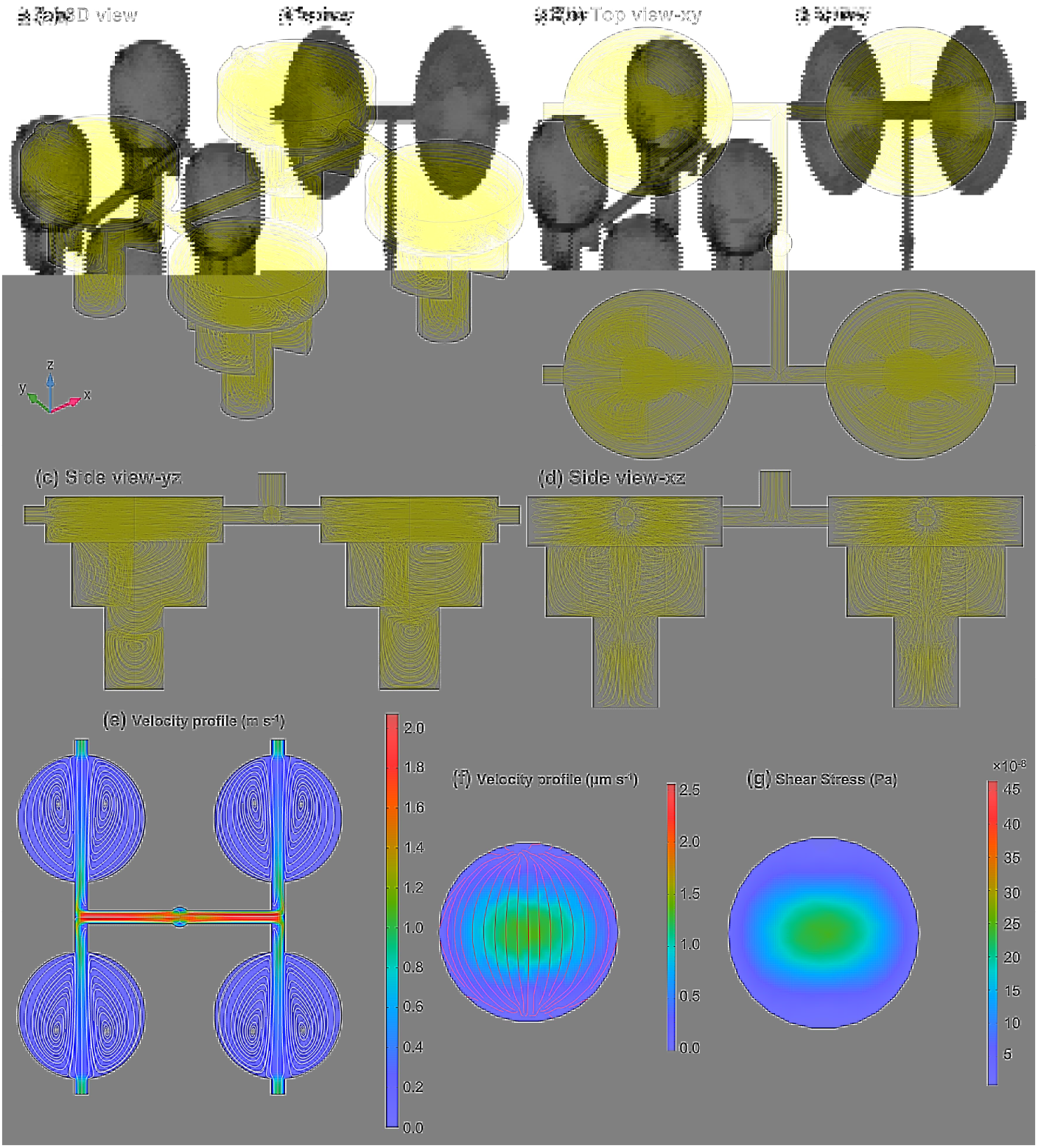
Quantitative simulation for IVES using COMSOL Multiphysics: (a) a 3D view of the IVES with gas (CO_2_) flow streamlines showing how gases distribute, (b) top view with gas flow streamlines, (c) and (d) the side views with gas flow streamlines. (e) The top-view of the velocity profile for the modeled gas presenting a uniform flow distribution among all four exposure chambers with a gentle velocity decrease, (f) the velocity profile at the close approximation to the surface where the cells were cultured, and (g) the shear stress profile at the location that the cells were cultured.

Internal gas diffusion inside a chamber was quantified with carbon dioxide (CO_2_), a product of combustion, as the gas of interest with an assumption of 1 mol L^−1^ at the inlet with normal diffusion in the air. Since the gas was uniformly distributed among all four exposed chambers (**Figure 3**), only one chamber was considered in the modeling (the gas flow at the inlet was assumed to be 7/4 mL s^−1^). The simulation was studied dependent on time, and the CO_2_ diffusion was modeled over 5 seconds, which was the time of exposure. A full cycle of smoke-fresh air was studied, and the cycle included one smoke exposure for 5 seconds (the initial concentration was set to zero; it was assumed that there was no CO_2_ in the chamber at the beginning) and one continuous fresh air exposure for 15 seconds (the initial concentration was taken from the final average volumetric concentration of the smoke exposure after 5 seconds resting based on the experimental set-up). To model the mass transport of CO_2_ in the chamber, Fick’s law was used as described below:

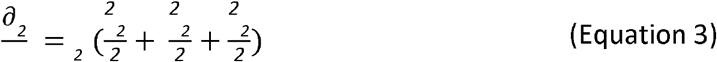

Where _2_is the carbon dioxide concentration in the smoke exposure chamber and_2_ (_2_ = 0.16 cm^2^ s^−1^)^[1]^ is the diffusion coefficient of CO_2_ molecules in the chamber.

The concentration distribution of CO_2_ in the chamber over 5 seconds of exposure is presented in **Figure 4a** with streamlines representing the concentration gradient in the chamber. After each smoke exposure, there was a 10-second resting time to allow the CO_2_ concentration to become uniform in the entire chamber. Therefore, the initial concentration for the fresh air exposure was calculated from the previous step and assumed to be uniform in the entire chamber, as seen in **Figure 4b**. **Figure 4b** also shows the rate of change and uniformity of the CO_2_ concentration in the chamber during fresh air exposure. **Figure 4c** and **Figure 4d** show the volumetric average CO_2_ concentration of the chamber and the average CO_2_ concentration at the outlet of the chamber, respectively. The simulation results confirmed that the CO_2_ concentration in the chamber reached zero after the first fresh air exposure, suggesting that the cells would experience a similar pattern for each cycle, thereby the size of the smoke exposure chamber was small enough to let a fast and uniform diffusion occur in the chamber. This means that the cells would experience the same gas concentration in each cycle.

**Figure 4:**
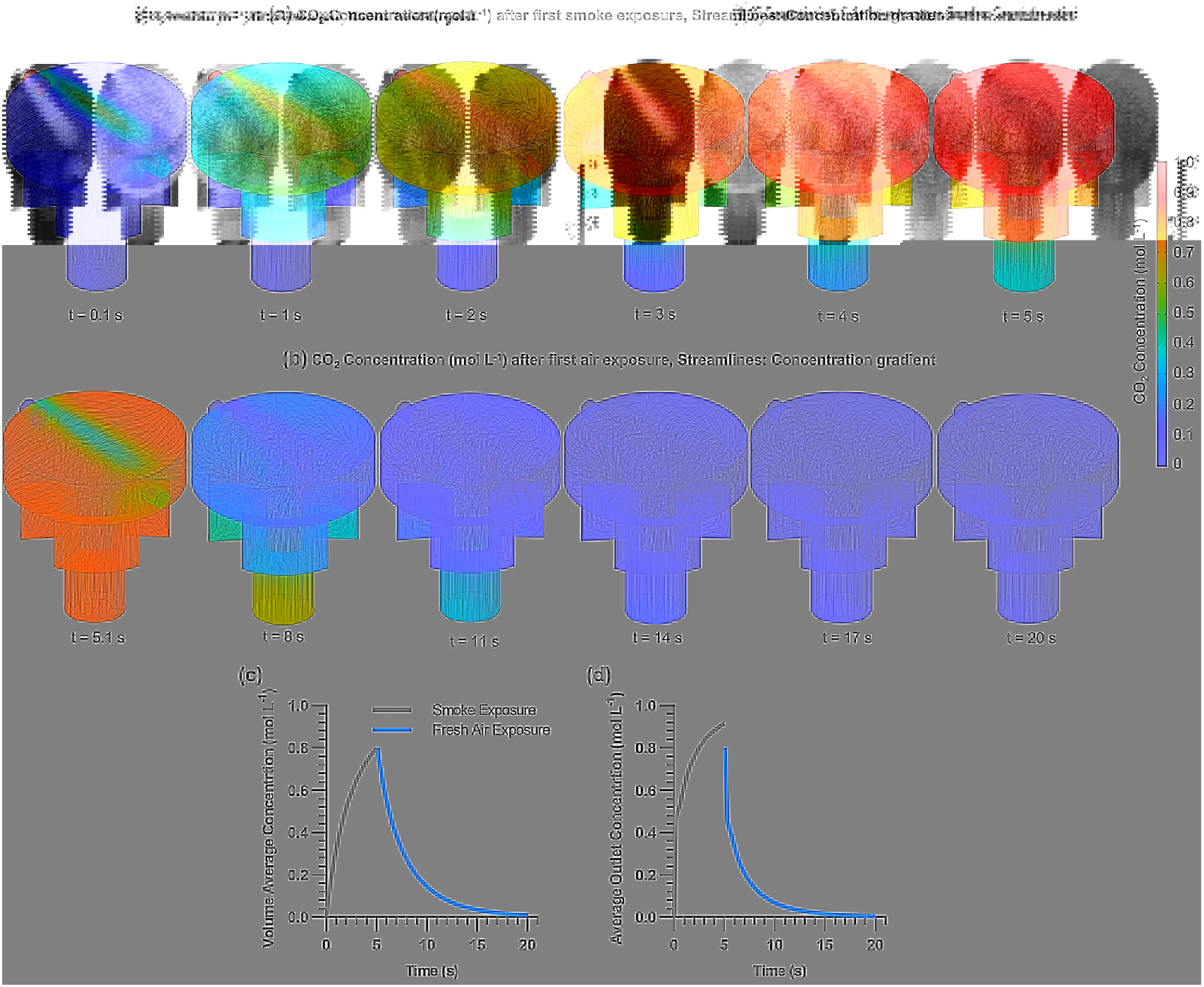
3D quantitative modeling results of the gas (CO_2_) concentration distribution in an IVES chamber: (a) real-time CO_2_ concentration distribution over 5 seconds exposure of smoke (initial CO_2_ concentration modeled at 1.0 mol L^−1^) showing a gentle gas distribution in the exposure chamber, (b) real-time CO_2_ concentration distribution over 5 seconds exposure of fresh air (the initial CO_2_ concentration was the final CO_2_ concentration in the exposure chamber from the previous smoke exposure and it was assumed that there was no CO_2_ in the fresh air exposed to the chamber), (c) volume average concentration of CO_2_ in the chamber for one smoke-fresh air cycle would lead to drop CO_2_ concentration in the chamber back to zero, and (d) average outlet concentration of CO_2_ after one smoke-fresh air cycle confirming that exposure kinetics were sufficient to reach a repeatable smoke/air exposure cycle (zero concentration at the outlet).

### Impact of *in vitro* whole cannabis smoke exposure on airway epithelial cell viability and barrier function

Following quantitative modeling, we next applied IVES for cannabis smoke exposure experiments with multiple biological readouts of relevance to epithelial cell biology. We measured transepithelial electrical resistance (TEER) before and after fresh, whole cannabis smoke exposure on primary human bronchial epithelial cells. We compared the change in TEER (Δ TEER) pre- and post-exposure between cultures that received room air or smoke. Our results suggest that individual donor cultures exposed to cannabis smoke in IVES experienced a decrease in epithelial barrier function as compared to air-exposed control (*p*<0.05) (**Figure 5A**). The decline in epithelial barrier function was not associated with any increase in LDH, a measure of cell membrane integrity and cell viability (**Figure 5B**) suggesting that cell cytotoxicity was minimally impacted by our model.

**Figure 5.**
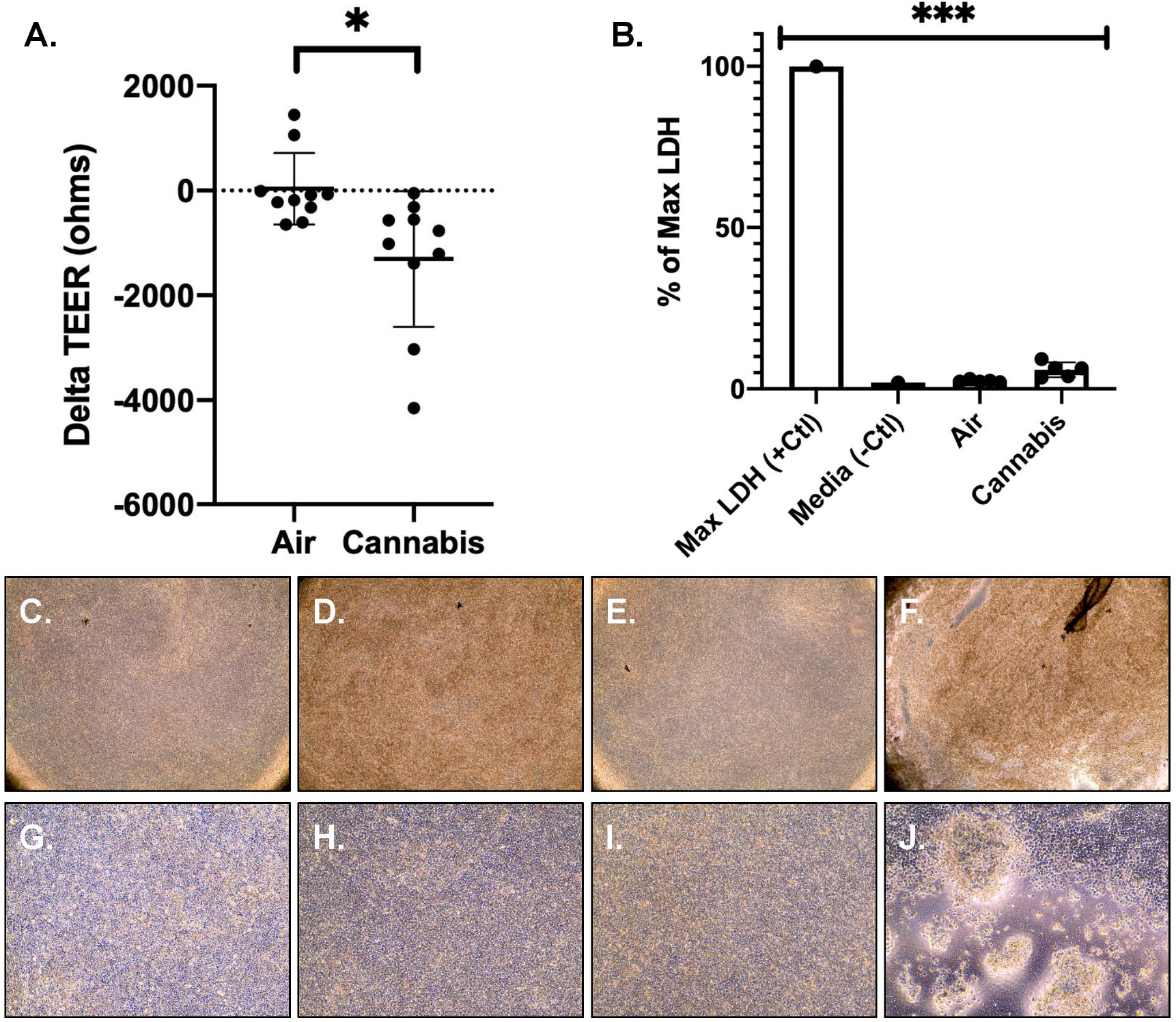
**(A)** shows the change in TEER from baseline after room air versus whole cannabis smoke exposure. Analyzed with paired t-test, p=0.029, n=10. **(B)** Lactate dehydrogenase expression as a proportion of maximal LDH release. Analyzed via ANOVA and Tukey’s post hoc test. Shows representative microscopy (40x) of **(C)** Transwell^®^ inserts with HBECs prior to room air exposure **(D)** Transwell^®^ with HBECs after room air exposure **(E)** Transwell^®^ with HBECs prior to whole cannabis smoke exposure **(F)** Transwell^®^ with HBECs after whole cannabis smoke exposure (product weight of 0.7g). Shows representative microscopy (100x) of **(G)** Transwell^®^ inserts with HBECs prior to room air exposure **(H)** Transwell^®^ with HBECs after room air exposure **(I)** Transwell^®^ with HBECs prior to whole cannabis smoke exposure **(J)** Transwell^®^ with HBECs after whole cannabis smoke exposure (product weight of 0.7g).

Following whole smoke exposure, cell cultures exhibited qualitative changes in morphology. Qualitative analyses revealed higher incidences of cell lifting, areas of patchiness and a circular shape of the cells relative to control. A representative microscopic image reflecting these notable changes is shown in **Figure 5**.

### Impact of *in vitro* whole cannabis smoke exposure on airway epithelial cell immune responses

IL-1 cytokine family members are elevated in the context of tobacco smoke exposure to airway epithelial cells and lung tissue (Churg et al., 2009; Kang et al., 2007; Pauwels et al., 2011). We therefore analyzed the differential expression of selected IL-1 cytokine family members - IL-1α, IL-1β, IL-18, IL-1Ra – in our model of cannabis exposure using IVES, as we have demonstrated significant overlap between airway epithelial cell responses to these two exposures in a submerged monolayer culture system (Aguiar et al., 2019). Trends for elevations in IL-1α (p=0.054), IL-1β (p=0.296) and IL-18 (p=0.064) were observed and a significant elevation of IL-1Ra (*p*<0.05) in five individual donor samples (n=5) following cannabis smoke exposure relative to room air. On a donor basis, IL-1α, IL-18 and IL-1Ra were elevated relative to control air in 5/5 samples (100%) whereas IL-1β was elevated in 4/5 samples (80%) (**Figure 6**).

**Figure 6.**
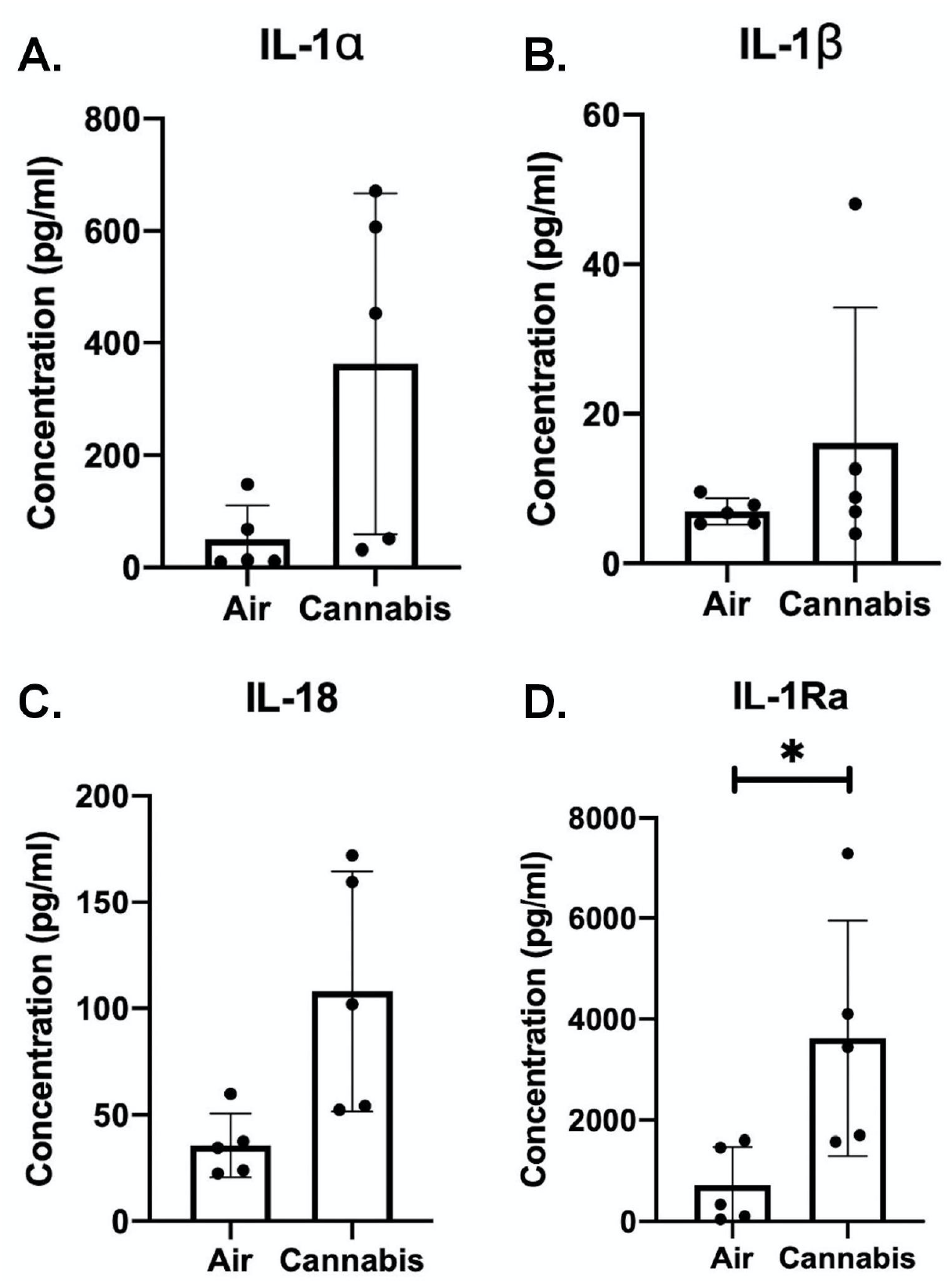
IL-1 cytokine family member quantification in apical washing of primary human airway epithelial cells exposed to air or whole cannabis smoke. **(A)** IL-1α, p=0.054 **(B)** IL-1β, p=0.296 **(C)** IL-18, p=0.064 **(D)** IL-1R antagonist, p<0.05 Analyzed via paired t-tests.

We have demonstrated that cannabis smoke extract suppresses CXCL10 and CCL5 in Calu-3 epithelial cells under submerged monolayer conditions (Aguiar et al., 2019). We therefore analyzed CXCL10 and CCL5 levels following cannabis smoke exposure in IVES with primary human airway epithelial cells. We observed no significant changes in both CXCL10 or CCL5 expression (**Figure 7**). On a donor basis 4/5 (80%) of donor samples displayed a suppression of CXCL10 (p=0.110) and 0/5 (0%) donor samples displayed a suppression of CCL5 relative to control air (p=0.252).

**Figure 7.**
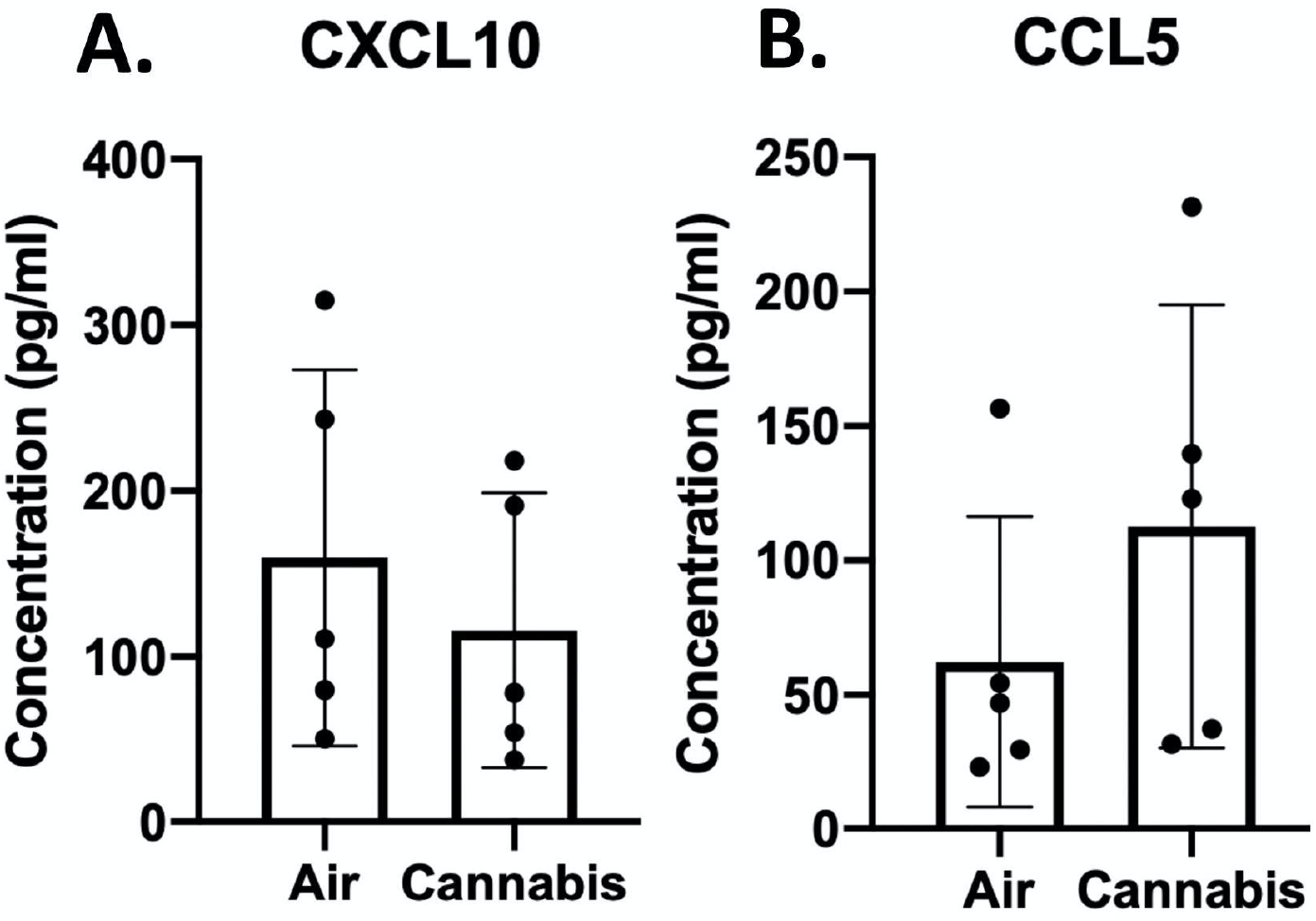
Antiviral cytokine quantification in apical washing of primary human airway epithelial cells exposed to air or whole cannabis smoke. **(A)** CXCL10, p=0.110 **(B)** CCL5, p=0.252 (n=5). Analyzed via paired t-tests.

### *In vitro* whole cannabis smoke exposure induces expression of genes involved in cellular detoxification

We have demonstrated that tobacco and cannabis smoke extract exposures with Calu-3 epithelial cells cultured under submerged monolayer conditions results in elevations in gene expression for CYP1A1 and CYP1B1, both which function as phase II detoxification enzymes (Aguiar et al., 2019). Additionally, Thioredoxin Interacting Protein (TXNIP) has been shown to inhibit the antioxidative effect of thioredoxin resulting in an accumulation of reactive oxygen species and cellular stress (Ji Cho et al., 2019; Junn et al., 2000). We therefore determined the gene expression level of CYP1A1, CYP1B1 and TXNIP in primary human airway epithelial cells following cannabis smoke exposure with IVES. We demonstrate that cannabis smoke exposure results in robust induction of both gene transcripts in 5/5 (100%) donor samples for CYP1A1 (*p*<0.001), 5/5 (100%) donor samples for CYP1B1 (*p*<0.05) and a suppression of TXNIP in 4/5 (80%) donor samples (*p*=0.058)(**Figure 8**).

**Figure 8.**
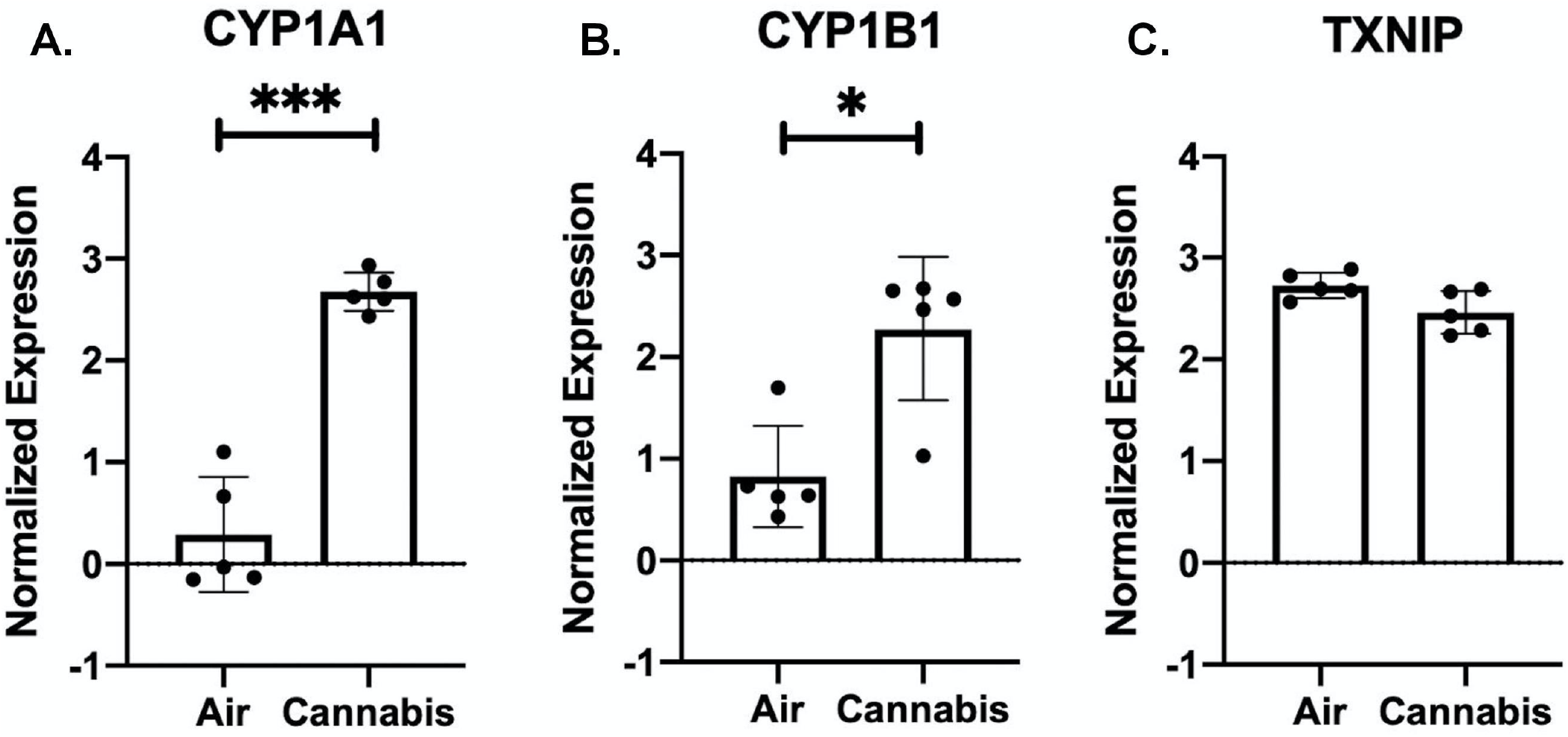
Oxidative stress genes expressed in primary human airway epithelial cells exposed to air or whole cannabis smoke. **(A)** CYP1A1, p< 0.001 **(B)** CYP1B1, p<0.05 **(C)** TXNIP, p=0.058 (n=5). Analyzed via paired t-tests.

## Discussion

The United States Center for Disease Control (CDC) declares that cigarette smoking is the leading preventable cause of death in the United States (CDCTobaccoFree, 2019). Adding to the pressing concern is the relatively new increase in global legalization and practice of smoking cannabis (Cerdá et al., 2017; Leyton, 2019; Volkow et al., 2014). Notably, evidence suggests that cannabis smoke consists of differential components relative to tobacco smoke, which may function as immunomodulatory agents (Ashton & Hancox, 2018; Kaplan et al., 2008; Klein, 2005). To determine shared or divergent consequences of cannabis smoke on lung health, relevant experimental data must be generated to inform policy and decision making at the personal and population level. Frequently used low cost *in vitro* models of smoke exposure may not adequately reflect the realities of smoke delivery *in situ*, while advanced smoke exposure model systems, more physiologically relevant, are often expensive and inaccessible for many research groups. To address these constraints and the need for accessible, relevant, *in vitro* experimental systems, we sought to develop and validate a novel smoke exposure model applicable to various modalities through the conception of IVES.

To capture the benefit of exposing cells directly to smoke while overcoming the shortcomings of the current smoke exposure systems, which are large, expensive, or lacking of adaptation to custom experimental set-ups (Amatngalim et al., 2015; de Bruijne et al., 2009; Herr et al., 2009; Lenz et al., 2009; Raredon et al., 2015), we have utilized 3D-printing technology to design and develop a versatile smoke exposure system that can be easily scaled up or down. Initially, the IVES was designed and fabricated as a single-chamber exposure unit. The purpose of this step was to optimize the size of the chamber considering various criteria such as basal liquid volume, accessibility to Transwell^®^ insert and the basal media, and the fluidic path of smoke exposure. Then, the final design was scaled up to the current IVES as proposed in this work (**Figure 1** and **Figure 2**). The fluidic dynamic modeling conducted in this study confirmed that the design of IVES and the exposure protocol were well-designed to expose cells in a repeatable fashion. **Figure 3** shows a uniform gas flow distribution in IVES with indirect gas exposure to cells, which was critical in creating a stress-free exposure to cells. Additionally, mass transfer results from the simulation revealed that the length of fresh air exposure was sufficient to zero CO_2_ concentration (**Figure 4**). This *in vitro* model can be easily redesigned for different applications and purposes. The size of IVES can be changed to fit for various sizes of Transwell^®^ inserts, or more exposure chambers can be integrated with IVES. Moreover, a fluidic path can be added to the basal side of the chambers to create a dynamic flow in the liquid compartment of the system. Overall, the proposed exposure system in this work is inexpensive, easy to use, easy-to-fabricate, and amenable to multiple experimental designs including smoke, vaping, co-exposures with pathogens.

The mucociliary escalator, secreted antimicrobial products and paracellular permeability mediated by intracellular junctions collectively serve to establish the barrier function between the external and internal environments (Ganesan et al., 2013). Human airways rely on the latter, tight and adherens junctions between cells at the apical border, to prevent inhaled pathogens and other insults from causing harm to the airways (Ganesan et al., 2013). In smokers, the epithelium is found to be dysfunctional and abnormally differentiated leaving a higher risk for viral, bacterial infection (Ganesan et al., 2013). To determine how IVES delivery of cannabis smoke impacted epithelial barrier function, TEER was measured in 10 individual donor primary HBECs prior to and following cannabis smoke exposure. Using IVES as a model for cannabis smoke exposure using primary HBECs, our data suggest that a compromised barrier may be observed in human cannabis smokers. The lungs rely on the formation and strict regulation of a mechanical barrier established by airway epithelial cells (Heijink et al., 2012). Previous studies have implicated oxidative stress brought on by cigarette smoke with the disruption of epithelial barrier function (Boardman et al., 2004). Other studies have found that cigarette smoke increases the permeability of human airways disrupting the balance between external fluids and macromolecules through altered regulation of multiple tight junction and adherens junction proteins (Mishra et al., 2016; Olivera et al., 2007; Tatsuta et al., 2019). Despite extensive studies of airway epithelial cell barrier function and tobacco smoke exposure, little data is available for cannabis smoke exposure. Using submerged monolayer cultures of Calu-3 cells, we were able to demonstrate that cannabis smoke condition media was able to reduce barrier function, as assessed by TEER, a response shared with tobacco smoke conditioned media (Aguiar et al., 2019). In the present study, we extended our published findings by interrogating how whole cannabis smoke impacted barrier function in primary human airway epithelial cells grown under ALI-conditions. Our data confirms that cannabis smoke, similar to tobacco smoke, is able to compromise barrier function. Furthermore, the data establish that monitoring epithelial cell barrier function measurements following exposures using IVES are possible.

Cytokines are signaling molecules crucial to the proper innate and adaptive immune function. They perform a host of essential duties ranging from mitigating viral, bacterial, fungal infections to signaling a cascade of other immunomodulatory agents responding to allergens in the air. Airway inflammation associated with changes in cytokines that regulate immune function is present in both cannabis and tobacco smokers resulting in clinical presentation of coughing, wheezing and the onset of asthma and COPD (Aldington et al., 2007; Costenbader & Karlson, 2006; Hancox et al., 2015; Joshi et al., 2014; Lee et al., 2012; Moir et al., 2008; Nagarkatti et al., 2009). Notably, the IL-1 family of cytokines has been implicated in acute inflammatory processes as well as linked to cytokine balance disruption in cigarette smokers (Nyunoya et al., 2014). IL-1 family cytokine expression at the protein level was assessed in our model following cannabis smoke exposure in five individual patient donor samples. Our experiments with five independent donors show trends for an increase in IL-1α, IL-1β, and IL-18 expression following cannabis smoke exposure relative to control, findings that are similar to observations made in other studies with tobacco smoke (Churg et al., 2009; Kang et al., 2007; Pauwels et al., 2011). These results indicate an inflammatory response induced by cannabis smoke characterized by IL-1 family cytokines, which may share downstream consequences with tobacco smoke that include neutrophilia driven by an IL-1R/IL-17 axis (Roos et al., 2015). We also observed a significant elevation of IL-1Ra following smoke exposure. Notably, other studies have found a suppression of IL-1Ra expression in tobacco smokers which works by antagonizing IL-1α and IL-1β (Shiels et al., 2014). In cannabis smoke however, IL-1Ra expression has been shown to be elevated owing to the unclear immunomodulatory features of cannabis active components (Melamede, 2005; Molina-Holgado et al., 2003).

Our published *in vitro* data using Calu-3 airway epithelial cells under submerged monolayer culture conditions suggest that cannabis smoke extract conditions media attenuates expression of antiviral pathways important in host defence in human airway epithelial cells (Aguiar et al., 2019; Huff et al., 2020). To explore the possibility that whole cannabis smoke exposure of primary human airway epithelial cells grown under ALI-culture conditions behaved similarly, we analyzed CXCL10 and CCL5 levels as previously performed (Aguiar et al., 2019; Huff et al., 2020). The current data shows negligible changes from the baseline of CXCL10 and CCL5 in primary HBECs. This suggests inherent differences in the model and/or use of primary cells at ALI when compared to cell-lines in submerged monolayers. Moreover, cytokines have been shown to not be expressed differentially following cigarette exposure alone; rather, they require viral/viral mimetic challenges prefacing expression (Eddleston et al., 2011; M. H. Hudy et al., 2010; M. H. Hudy et al., 2013). It will be relevant for future studies to induce CXCL10/CCL5 through viral/viral mimetic challenges to assess the impact of cannabis smoke on these antiviral cytokines.

Previously, we have shown that aryl hydrocarbon receptor induced genes associated with oxidative stress, CYP1A1 and CYP1B1, are significantly induced in Calu-3 cells exposed to cannabis and tobacco smoke extracts (Aguiar et al., 2019). Similarly, we have shown that TXNIP was reduced (Aguiar et al., 2019). Other studies have shown similar results of increased oxidative stress as indicated by dysregulated expression of these same genes in various types of smoke exposure such as tobacco, polycyclic aromatic hydrocarbons, incense smoke and cannabis smoke (Al-Arifi et al., 2012; Hussain et al., 2014; Roth et al., 2000). Consequently, we analyzed the expression of CYP1A1, CYP1B1 and TXNIP in primary human epithelial cells exposed to cannabis smoke using IVES. We chose these particular genes because of their robust induction in our Calu-3 cell model with cannabis smoke conditioned media. A significant elevation was observed in CYP1A1 and CYP1B1 while a negative trend was observed in TXNIP following smoke exposure, validating IVES for modeling molecular changes in human airway epithelial cells exposed to whole cannabis smoke.

The aim of our study has been to validate and introduce an open-access, disposable, easy to use method for whole smoke exposure to airway epithelial cells grown on Transwell^®^ inserts. The dosing parameters used in our study served as a means to reliably evoke a cannabis induced effect. Herein, we validate IVES for equal distribution of smoke across the four exposure chambers, equal distribution of smoke pressure over the base of the Transwell^®^ insert, minimal shear stress across the growth area, a capacity for the human airway epithelial cells to respond to smoke at a cellular and molecular level. Future studies can expand applications for the validated IVES to explore different concentrations, durations of smoke, period of smoke, co-exposure with pathogens, or introduction of vaping technologies. Additional experiments could be performed to define a no-observed effect level for studies with toxicology focus.

In this study, we have outlined and validated the developmental parameters, flow simulation and streamlined the exposure protocol to measure epithelial barrier function, cytokine expression and gene expression. We have applied it to an exposure study assessing the impact of cannabis smoke exposure of primary HBECs. IVES shows stark similarities between existing models and promises the ability to generate needed relevant data to inform public health policy and individual user practices.

